# pSONIC: Ploidy-aware Syntenic Orthologous Networks Identified via Collinearity

**DOI:** 10.1101/2021.02.18.431864

**Authors:** Justin L Conover, Joel Sharbrough, Jonathan F Wendel

**Author notes:** Corresponding author: Jonathan F Wendel, Mailing address: Department of EEOB, 251 Bessey Hall, 2200 Osborn Dr, Ames, IA 50011.

## Abstract

With the rapid rise in availability of high-quality genomes for closely related species, methods for orthology inference that incorporate synteny are increasingly useful. Polyploidy perturbs the 1:1 expected frequencies of orthologs between two species, complicating the identification of orthologs. Here we present a method of ortholog inference, Ploidy-aware Syntenic Orthologous Networks Identified via Collinearity (pSONIC). We demonstrate the utility of pSONIC using four species in the cotton tribe (*Gossypieae*), including one allopolyploid, and place between 75-90% of genes from each species into nearly 32,000 orthologous groups, 97% of which consist of at most singletons or tandemly duplicated genes -- 58.8% more than comparable methods that do not incorporate synteny. We show that 99% of singleton gene groups follow the expected tree topology, and that our ploidy-aware algorithm recovers 97.5% identical groups when compared to splitting the allopolyploid into its two respective subgenomes, treating each as separate “species”.

## INTRODUCTION

The recent explosion in high-quality genome assemblies has increased the opportunity to investigate biological questions using a comparative genomics framework. An essential first step in many applications is inference of a high-confidence set of orthologs in the genomes under study. Methods for inferring orthologs are broadly based on sequence similarity, either through the construction of phylogenetic trees or through clustering of sequence similarity scores. There has been considerable progress in developing methods that curate a genome-wide set of orthologs for distantly-related genomes (Trachana *et al.* 2011; Emms and Kelly 2020), prioritizing flexibility for use on species with fragmented genome assemblies or even transcriptome assemblies, e.g. Inparanoid (O’Brien *et al.* 2005), OrthoMCL (Li *et al.* 2003), and OrthoFinder (Emms and Kelly 2015, 2019). As genomes for closely-related species become more prevalent, however, methods designed for deep-phylogenetic identification are less than optimal, as new methods can leverage conserved gene order across closely-related species (i.e. synteny) as powerful evidence for orthology. Two closely related species have largely collinear genomes, barring chromosomal rearrangements or small-scale gene loss or gain events (e.g. via transposition) that break up blocks of collinear genes (Dehal and Boore 2005). Programs have been developed to identify these collinear blocks (e.g. MCScanX (Wang *et al.* 2012) and CoGe (Lyons *et al.* 2008)) but these methods are restricted to pairwise comparisons (MCScanX) or comparisons among three genomes (CoGe), and no method for genome-wide detection of orthologs across multiple species has yet incorporated the powerful evidence of orthology provided by synteny.

A biological feature that can complicate orthology inference is whole genome multiplication (polyploidy), which is widespread throughout the tree of life (Van de Peer *et al.* 2017; Li *et al.* 2018), especially in plants (Jiao *et al.* 2011; One Thousand Plant Transcriptomes Initiative 2019). In the case of ancient polyploids, extensive gene deletion and chromosome rearrangement (Wendel 2015) often obscures the expected number of gene copies that should be present in a genome and complicates pairwise genome alignments, syntenic block detection, and gene tree - species tree reconciliation. Differences in ancestral ploidy levels have not been integrated into existing programs for detecting orthologs, although this is essential for obtaining accurate estimates of orthogroup completeness.

Here, we present a new method of ortholog inference, Ploidy-aware Syntenic Orthologous Networks Identified via Collinearity (pSONIC), which uses pairwise collinearity blocks from multiple species inferred via MCScanX, along with a high-confidence set of singleton orthologs identified through OrthoFinder, to curate a genome-wide set of syntenic orthologs. As part of pSONIC’s inference, we developed a ploidy-aware algorithm to identify collinear blocks originating from both speciation and duplication events. To evaluate pSONIC’s performance in a system with a complex history of duplication and speciation, we tested pSONIC on four species in the cotton tribe (*Gossypieae*), including one allopolyploid, its two closest diploid progenitors, and a phylogenetic outgroup. Our method assigned between 75-90% of all genes into orthogroups, and when compared to OrthoFinder, identifies 40% more single-copy orthogroups (97% of which exhibit gene tree topologies consistent with the species relationships within the *Gossypieae*) and 33% more orthogroups that contain only tandemly duplicated genes from each species. To demonstrate the effectiveness of our ploidy-aware algorithm, we show that, unlike OrthoFinder, splitting the tetraploid genome into its respective genomes has little effect on our final set of orthogroups.

## METHODS

Required input for pSONIC includes the list of orthogroups inferred from OrthoFinder (-og flag) and the files containing the list of collinearity groups and tandemly duplicated genes from MCScanX (default parameters). To reduce the memory requirements of MCScanX and pSONIC, gene names are converted to the style of OrthoFinder (this can be done using the --translate_gff flag in the pSONIC program). Additionally, an optional file providing the relative degree of ploidy increase for each species can be used for analyses in which a polyploidy event has occurred along the phylogeny of the species in the analysis. In the example below from the cotton tribe, we run two analyses to demonstrate this: one in which the subgenomes of allopolyploid *Gossypium hirsutum* (2n = 4x) is run normally (i.e. the relative ploidy is 2 compared to all other species in the analysis), and a second in which the genome has been split into its respective subgenomes, with each treated as a separate “species”. This feature allows the possibility of syntenic analysis of genomes with varying complexities of ploidy histories, including those where clear partitioning into subgenomes is not possible.

The pSONIC pipeline proceeds in four basic steps. First, OrthoFinder results are parsed to find “tethers” -- that is, orthogroups in which at least two species have fewer than or equal to the number of “gene sets” expected from relative ploidy levels. Here, we define “gene sets” to include a gene and all immediately neighboring tandemly duplicated genes (as determined by MCScanX). Using “gene sets” instead of singleton genes dramatically increases the number of tethers that can be used in steps two and three without creating spurious cases of inferred collinearity (Table 1). For any orthogroup in which some, but not all, species contain more gene sets than expected by ploidy, genes from these species are excluded while genes from all other species are included in downstream steps. Our method also permits inference of orthology when specific orthologs are missing due to, for example, gene loss following polyploidy.

**Table 1:**
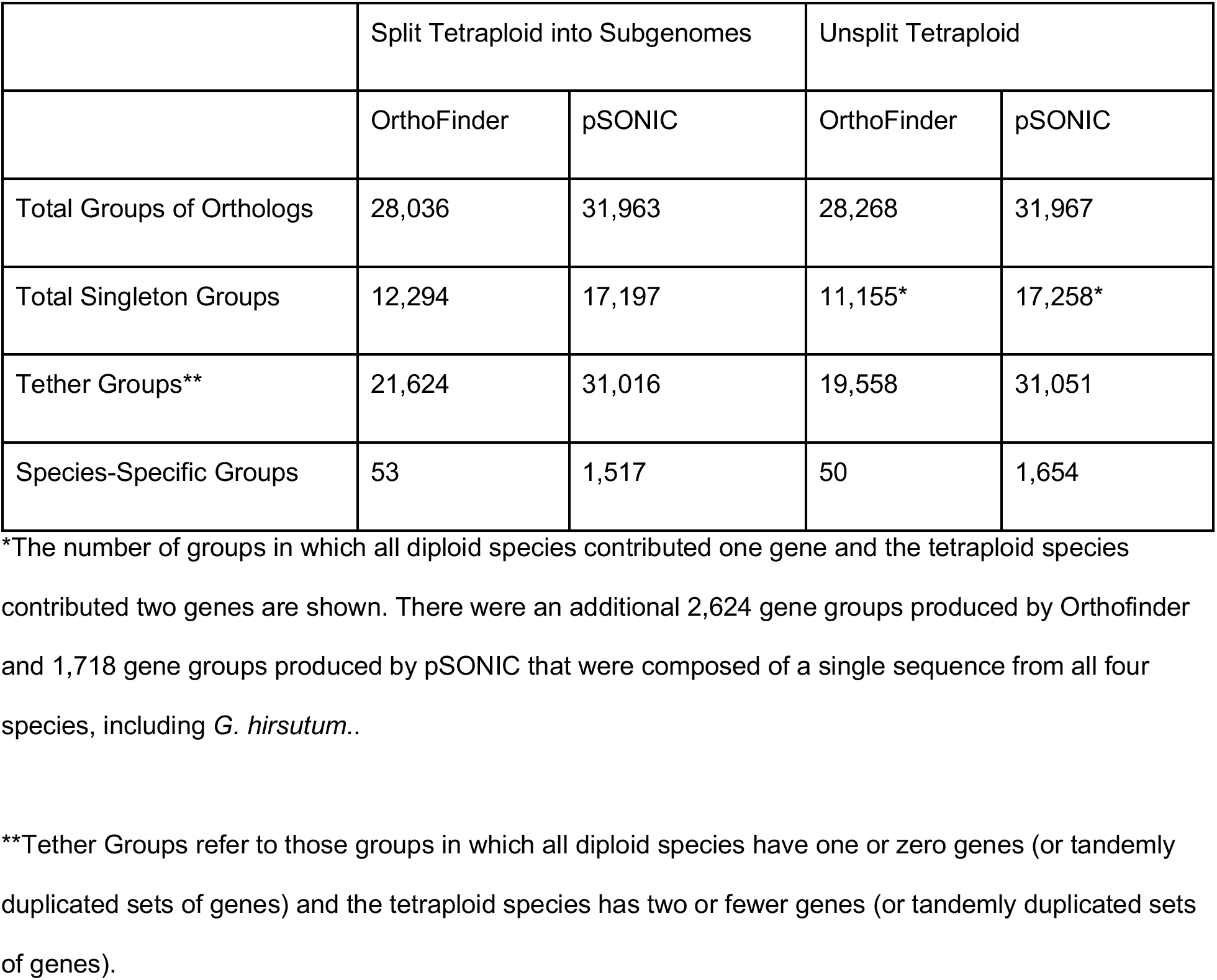
Summary Statistics of OrthoFinder versus pSONIC

**Table 2:**
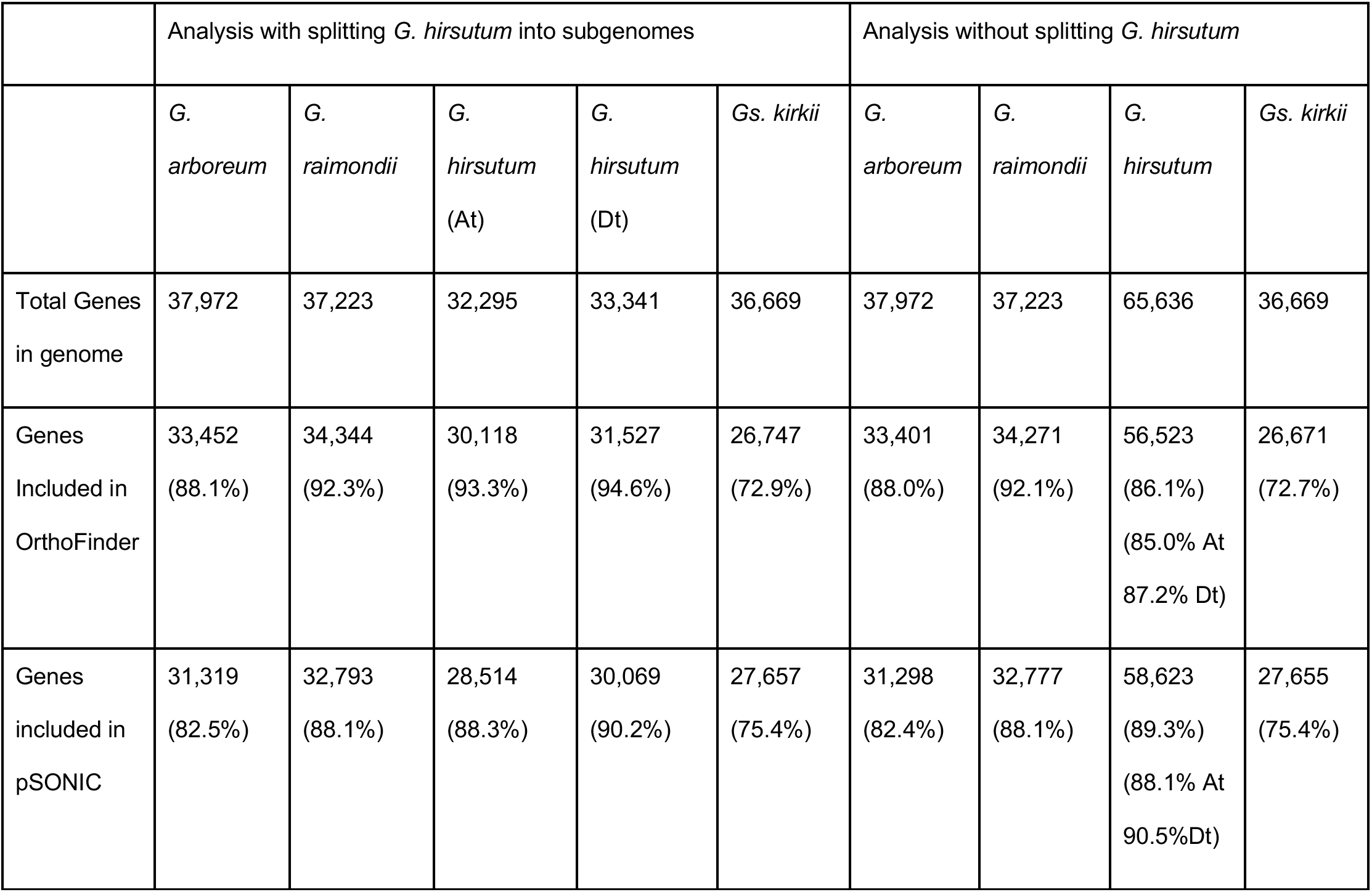
Performance of OrthoFinder and pSONIC When Splitting Tetraploid Genomes into Subgenomes

The second step of the pSONIC pipeline is to parse the output from MCScanX to find syntenic blocks that correspond to those present in the most recent common ancestor of all species in the analysis. Ancient duplication events create syntenic blocks (Figure 1), which MCScanX will identify as long as the genes in this block have sufficient protein sequence similarity. For each syntenic block, each pair of genes along the block is compared to the set of tethers described in Step 1. Each gene pair is given one of three classifications: (A) “Pass”; (B) “No Call”; or (C) “Not Pass”. The decision tree leading to this classification is described in Figure 2. Syntenic blocks that contain fewer than two gene pairs with “Pass” scores, or that contain more “Not Pass” scores than “Pass” scores, are discarded and removed from further analysis.

**Figure 1:**
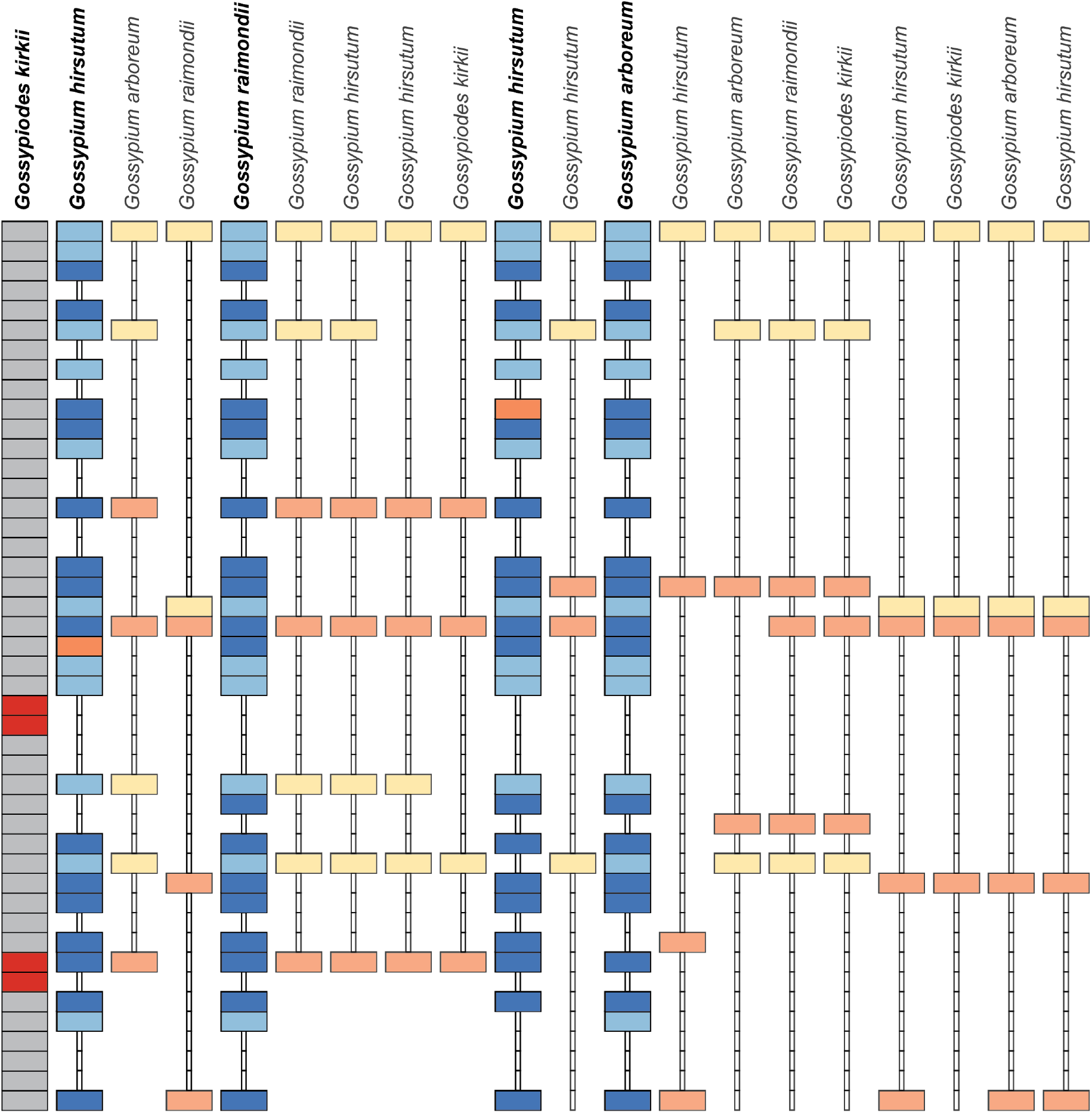
Syntenic blocks inferred by MCScanX mapped onto a single chromosomal region make for complicated multi-species inference of syntenic orthology. A sample of collinear blocks identified by MCScanX (one per column) is shown aligned to a segment of chromosome KI24 in *G. kirkii* (dark grey). Due to ancient whole genome duplication events, many syntenic blocks may map onto the same chromosomal region of a reference genome. Genes that are tandemly duplicated in the reference genome are shown in red. We classify each gene pair in each collinear block into one of three groups: Pass (dark blue) if both genes are in the same tether set; Not Pass (orange) if only one of the two genes is in a tether set, but the other gene is absent; and No Score (light blue/yellow) if neither gene is in a tether set. Light blue genes are included in the list of edges because the collinear group they belong to had more than two “Pass” scores, and more Pass scores than “Not Pass” scores (groups indicated by species names above in bold).

**Figure 2:**
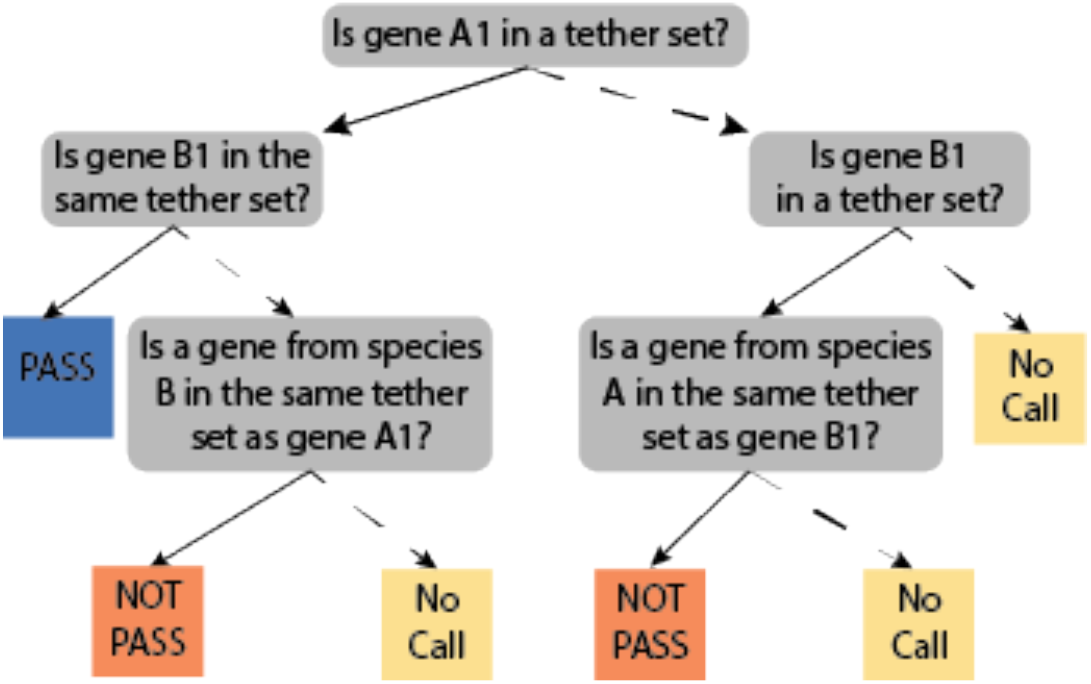
Decision Tree for Scoring Gene Pairs. Consider the first gene pair along a collinear block consisting of genes A1 and B1 from species A and B, respectively. pSONIC scores this gene pair based on a set of questions. Starting from the top-most question, if the answer is “yes”, follow the solid line; if the answer is “no”, follow the dashed line. Briefly, if both genes A1 and B1 are in the same tether set, they are classified as “Pass”, and if they are in different tether sets, they are classified as “Not Pass”. If neither gene is found in any tether set, they are classified as “No Call”. Finally, if the tether set that contains one of the genes (e.g. A1) also contains a gene from the other species (e.g. species B) that is not the gene pair in question (i.e. B1), then that gene pair is classified as “Not Pass”; however, if that tether set does not contain any genes from the other species (e.g. species B), then the gene pair is classified as “No Call”. Importantly, the above tree results in the exact same output regardless of which gene in the pair is considered A1. In the case of scoring a collinear block between two regions of the same tetraploid genome, species A and species B are the same, and the same decision tree is used.

Third, for those blocks that pass the filtering in Step 2, the ends of the collinear block are trimmed in the following way (Figure 3): reading from the end of the collinear block, if the first gene pair that did not receive a “No Call” designation is a “Not Pass” tether, all gene pairs up to and including that “Not Pass” gene pair are discarded (Figure 3B). Additionally, for the first six gene pairs that received a “Pass” or “Not Pass” score, if three or more of the scores are “Not Pass”, the end-most gene pair is trimmed from the block sequentially until this criteria is met (Figure 3C). Finally, to prevent two neighboring syntenic blocks from being incorrectly condensed into the same block, we also split any block that had three consecutive “Not Pass” scores without an intervening “Pass” score (Figure 3D). The ends of these two newly-created blocks are then re-trimmed as described above. These filtering procedures are repeated recursively until all criteria are fulfilled. If any block post-filtering contains fewer than five genes, that block is removed from downstream analyses. We implemented this filter because we found that towards the ends of some collinear blocks, there were several successive gene pairs that received “Not Pass” scores even though the block collectively received many more “Pass” scores, and adding this filter greatly increased the number of resulting tethered groups.

**Figure 3:**
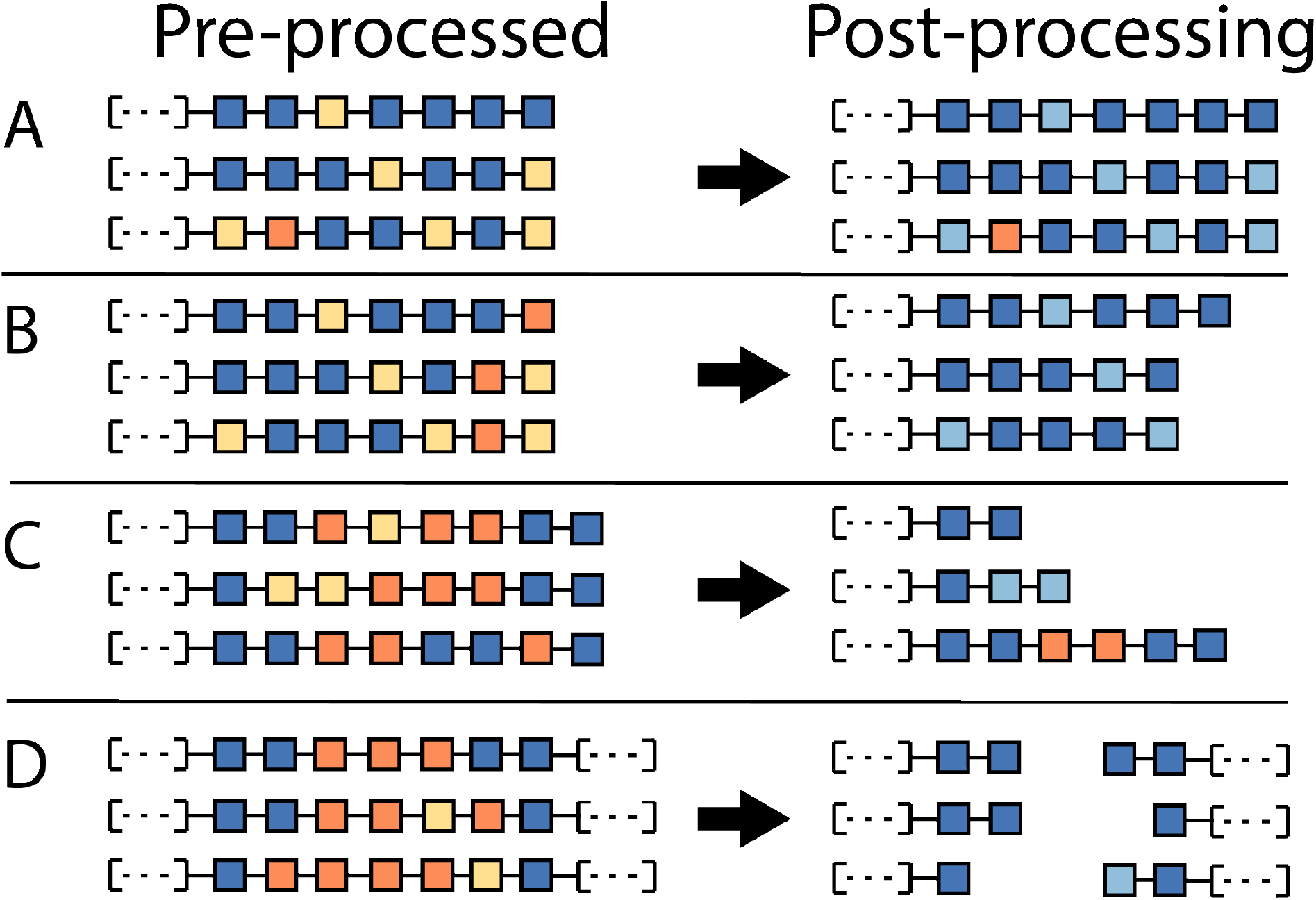
Syntenic block trimming using tethered genes improves quality of syntenic orthology inferences. For collinear blocks that have at least two “Pass” scores and more “Pass” scores than “Not Pass”, pSONIC trims the edges of the blocks to ensure proper cutoff placement of the block ends. **A.)** If the end of the collinear block contains a “Pass” score, or if no “Not Pass” gene pairs are more distal than the first “Pass” gene pair, then all gene pairs are retained and used as edges in the graph construction step of pSONIC. **B.)** If the end-most gene pair received a “Not Pass” score, then it is removed from the collinear block. Likewise, if no gene pair received a “Pass” score before the endmost “Not Pass” gene pair, then all gene pairs up to and including that “Not Pass” gene pair is removed from the collinear block. **C.)** For the first six gene pairs that receive either a “Pass” or “Not Pass” score, if the number of “Not Pass” scores is three or more, the endmost gene pair is removed from the collinear block. This process is repeated until the number of “Pass” gene pairs outnumber the “Not Pass” gene pairs. As depicted in the third row, some “Not Pass” gene pairs could still pass after this filter is applied. **D.)** If there is an internal segment of the collinear block that contains three or more “Not Pass” scores without any intervening “Pass” scores, the block is split into two, and the ends of the new blocks are trimmed using processes described in B and C. Each collinear block must pass all filters to be used in downstream steps.

Finally, all collinear blocks that pass Step 3 are assembled into a set of syntenic genes across all species. To do this, we first construct an empty graph where the vertices include all genes from all species. We then treat each pair of genes along every collinear block as edges of this graph. Tandem duplicates represent a specific case of syntenic orthology, and where present, are also included as edges in the graph. This graph is then decomposed into subgraphs, with each subgraph of more than two vertices representing a syntenic group of orthologs. Thus, by synthesizing pairwise gene-order collinearity tracts into multiple-species collinearity subgraphs using amino acid sequence similarity, we are able to infer genome-wide orthologs in extensively duplicated genomes and across multiple species, simultaneously.

## RESULTS AND DISCUSSION

### pSONIC Identifies Single-Copy Orthogroups with High Resolution

To show the utility of pSONIC, we created a genome-wide list of orthologs for genomes from four species in the cotton tribe (*Gossypieae*); specifically, allopolyploid *Gossypium hirsutum (Saski et al. 2017)*, two model progenitors of the polyploid (*G. raimondii (Paterson et al. 2012)* and *G. arboreum (Du et al. 2018)*), and an outgroup, *Gossypioides kirkii (Udall et al. 2019).* All protein sequences and gff files were downloaded from CottonGen (Yu *et al.* 2014) and only the primary isoforms of proteins located on the annotated chromosomes of each species were used in this analysis. All original files used are provided in the GitHub repository as a test data set, and BLAST files to run MCScanX can be found in File S1. While there is a history of complicated polyploidization events in the family (Malvaceae) to which cotton belongs (Conover *et al.* 2019), only one neoallopolyploidy event is included among the species chosen, making this an ideal system to demonstrate the utility and flexibility of pSONIC.

We first decided to split the subgenomes of allopolyploid *G. hirsutum* into its respective subgenomes, treating them as separate “species”. From this split input, OrthoFinder produced 28,036 orthogroups, 21,624 of which were classified as “tethers”, and 12,294 that contained exactly one gene sequence from each species (i.e., single copy orthogroups). MCScanX initially identified 20,392 collinear blocks between the five species, but only 1,833 passed the filtering criteria of pSONIC, highlighting the complex history of polyploidy in this tribe. After trimming and splitting these blocks, pSONIC assembled the remaining 238,452 edges into 31,963 groups of orthologs (Table 1). Of these, 17,197 contained exactly one gene from each species, and 31,016 groups contained at most one gene set (i.e. singleton gene or one tandemly duplicated set of genes). This 40% increase in singleton groups compared to OrthoFinder demonstrates a remarkable improvement in resolution of gene composition, and demonstrates the usefulness of pSONIC for analyses containing only diploid species.

To evaluate the quality of the orthologous relationships inferred by pSONIC, we quantified the extent to which single-copy gene groups reflected the phylogenetic history for these four well-differentiated species. We aligned CDS sequences using MAFFT v 7.407 (Katoh and Standley 2013), selected models of evolution using jModelTest v2.1.10 (Darriba *et al.* 2012), inferred gene trees using PhyML v20130103 (Guindon and Gascuel 2003), and compared tree topology for each of the single-copy genes identified by pSONIC to the known species relationships. Of the 17,197 single-copy genes, we found that 17,057 (99.2%) exhibited a tree topology consistent with the species tree. We also used *Gs. kirkii* to test root placement on the 17,057 topologically consistent gene trees and found that the root was between the A and D lineages in 15,950 (93.5%) gene trees (File S2). Together, these phylogenetic results indicate that the gene sets inferred by pSONIC are highly likely to be true orthologs.

### Efficacy of the Ploidy-Aware Algorithm

We also ran pSONIC while treating *G. hirsutum* as a single species instead of treating each subgenome as separate species to show the utility of the ploidy-aware algorithm. Interestingly, OrthoFinder placed fewer genes from all species into orthogroups, including 5,122 fewer genes from *G. hirsutum* and identified fewer singleton and tether groups than when the tetraploid genome was split *a priori* (Table 1). However, the poorer performance of OrthoFinder had a negligible effect on the number of genes from each species placed in orthogroups by pSONIC. Specifically, 40 more genes from *G. hirsutum* were placed into orthogroups in our ploidy-aware algorithm, and only 21, 16, and 2 fewer genes from *G. arboreum, G. raimondii*, and *Gs. kirkii* were included, respectively. pSONIC was able to identify 54.7% more (17,258 versus 11,155) singleton orthologous groups (i.e. groups in which the tetraploid had two genes and all diploids had one gene) and 58.7% more (31,051 versus 19,558) tether groups (i.e. groups in which the tetraploid had two or fewer genes or tandemly duplicated gene sets, while each diploid had at most one gene or tandemly duplicated gene set) than OrthoFinder.

When we compare the syntenic orthologous groups produced by the ploidy-aware algorithm to splitting the polyploid *a priori*, the results are largely identical. The two approaches produced 31,101 groups with identical gene membership. The ploidy-aware method identified 27 groups in which no genes were placed in the *a priori* split groups, while the *a priori* split method produced 52 groups in which no genes were placed in the ploidy-aware groups. There was a small proportion (~2.5%) of groups in which gene membership overlapped but was not identical across the two methods. The 810 groups recovered from the *a priori* split method that overlapped non-identically with 839 groups recovered from the ploidy-aware method formed 691 combined gene groups. Of these 691 overlapping gene groups, we found 570 (82.5%) in which the two methods directly conflicted about which genes from a given species were to be included, and 121 (17.5%) groups that did not disagree with respect to the genes from any given species, but included genes from individual species that were recovered by one method but not the other. In sum, the ploidy-aware algorithm agreed with the *a priori* ploidy determined set of genes in over 98.2% of cases, indicating that *a priori* splitting of polyploid subgenomes is not strictly necessary.

### Output Files

pSONIC produces several output files that describe the orthogroups, including the number of genes from each species included in each orthogroup, how many gene sets (i.e. sets of tandemly duplicated genes) from each species are included in each orthogroup, statistics from every collinear group (e.g. block size, how many “Pass” vs “No Pass” scores, etc), how the ends of every collinear block were trimmed and/or split, and which collinear groups were used in the final step of creating the final set of orthologs. Details about these individual files are explained in full in the README file in the Github repository.

### Data Availability

pSONIC is a program written in Python (written and tested on Python v3.7.7) and is freely available on GitHub (https://github.com/conJUSTover/pSONIC). Test data for running the program are provided on GitHub. Supplemental files available at FigShare. File S1 contains blast scores for both analyses of MCScanX and OrthoFinder. File S2 contains alignments, model selection, gene trees, and a summary file of all phylogenetic trees for over 17,000 singleton orthogroups identified by pSONIC.

## ACKNOWLEDGEMENTS

We thank the Iowa State University ResearchIT Unit for computational support. This work was supported by funding from NSF-Plant Genome Research Program (JS and JFW) and Cotton Inc. (JFW and JLC). This work utilized resources from the University of Colorado Boulder Research Computing Group, which is supported by the National Science Foundation (awards ACI-1532235 and ACI-1532236), the University of Colorado Boulder, and Colorado State University.

